# Fgf3 and Fgf10a regulate neuronal fasciculation through Schwann cell proliferation and infiltration in zebrafish posterior lateral line

**DOI:** 10.64898/2026.04.05.716528

**Authors:** Hao Jie Wong, Takaaki Matsui, Yasumasa Bessho, Ryutaro Akiyama

## Abstract

**Background:** During development, axons are organized into bundles, a process known as axonal fasciculation. The zebrafish lateral line nerve has been used as a model to study axonal fasciculation; however, the underlying mechanisms are not yet fully understood. Although Fgf3 and Fgf10a are well known to regulate the migration of the lateral line primordium along which the lateral line nerve projects, their roles in the organization of the lateral line nerve itself have not been clarified.

**Results:** *fgf3,10a* double mutants exhibited lateral line axonal defasciculation accompanied by an increased number of Schwann cells. Live imaging revealed a marked increase in Schwann cell proliferation and demonstrated that newly divided Schwann cells migrate along axons and infiltrate interaxonal spaces, thereby expanding these spaces and disrupting axonal fasciculation. Pharmacological manipulations further implicated a contribution of Nrg1-ErbB signaling to this phenotype.

**Conclusions:** Our findings suggest that Fgf3 and Fgf10a are required to restrict Schwann cell proliferation and infiltration, thereby ensuring axonal fasciculation during lateral line development.

## 1 INTRODUCTION

The zebrafish lateral line is a model system for studying the development of sensory organs. The posterior lateral line (pll) is a network of mechanosensory organs distributed along the body’s surface. These organs, called neuromasts, contain clusters of specialized sensory hair cells that detect water movement, enabling the fish to perform vital functions such as prey detection, predator avoidance, and social communication^1–4^. Because the lateral line develops externally along the body surface, it provides an excellent system for spatiotemporal analysis of the coordinated development of multiple cell types that constitute a sensory organ, including sensory cells, neurons, and glial cells.

The development of the pll begins at 20-22 hpf, during which the primordium migrates posteriorly along the horizontal myoseptum toward the caudal fin^5^. As the primordium migrates, its cells reorganize sequentially to form protoneuromasts and periodically deposit neuromasts along its path (22-42 hpf). Following the departure of the primordium, axons extend from the pll ganglion and are guided by cues derived from the migrating primordium^5–9^. Schwann cells (a type of glial cells in the peripheral nervous system) then divide and migrate along these extending axons to ensheathe them^10^. Subsequently, primordium migration is completed by 42 hpf ^11^, and the deposited neuromasts differentiate into supporting cells and hair cells, where axons innervate to form complete functional mechanosensory units capable of perceiving environmental stimuli^12^.

Fibroblast growth factors (Fgfs) are secreted signaling molecules that play critical roles in development^13^. Among them, Fgf3 and Fgf10a are considered key regulators of zebrafish posterior lateral line (pll) development. During pll development, premigratory and migrating primordium express Fgf3 and Fgf10a, and the morphogenesis and migration of the pll primordium are regulated by these factors through the MAPK (mitogen-activated protein kinase) signaling pathway^14–16^. Loss of function of Fgf3 and Fgf10a disrupts the epithelialization and compartmentalization of protoneuromasts, impairing rosette assembly and subsequently affecting primordium migration^14,16^.

Prior to the primordium migration, the pll ganglion is positioned immediately anterior to the primordium, placing lateral line neurons in close spatial proximity to *fgf3*- and *fgf10a*-expressing tissues. During migration, the primordium remains closely associated with extending lateral line axons and Schwann cells, which together establish the sensory circuitry of the lateral line system. Given that Fgfs are diffusible signaling molecules, these spatial and temporal relationships raise the possibility that Fgf3 and Fgf10a act not only on the primordium itself but also on neighboring lateral line neurons. Despite the well-established roles of Fgf3 and Fgf10a in primordium morphogenesis and migration, it remains unclear whether these factors regulate lateral line neuron development.

In this study, we revisited posterior lateral line (pll) development in zebrafish lacking *fgf3* and *fgf10a* to investigate their roles during lateral line neuronal development. We show that *fgf3* and *fgf10a* are required for proper axonal fasciculation in the pll, and that loss of these factors results in disorganized axons. Live imaging revealed excessive Schwann cell proliferation in *fgf3,10a* double mutants, with newly divided daughter Schwann cells infiltrating interaxonal spaces, leading to axonal defasciculation. Pharmacological suppression of Schwann cell proliferation *via* inhibition of ErbB signaling rescued the axonal bundling defects, indicating that aberrant Schwann cell proliferation underlies the neuronal phenotype. Furthermore, we found that Nrg1 expression is upregulated in the pll ganglion of *fgf3,10a* mutants, and that neuronal overexpression of Nrg1 phenocopied the Schwann cell and axonal defects. Together, these findings identify a previously unrecognized role of Fgf3 and Fgf10a in coordinating neuronal morphology through the regulation of Schwann cell proliferation during sensory system development.

## 2 RESULTS

### 2.1 Characterization of *fgf3,10a* double mutants generated through CRISPR-Cas9 technology

To assess the roles of Fgf3 and Fgf10a in lateral line nerve development in zebrafish embryos, we generated the loss-of-function mutants by targeting the first coding exon as shown in Figure 1A. Mutations in the F1 progeny were identified, resulting in 13-base pair (bp) and 7 bp deletions in *fgf3* and *fgf10a*, respectively. These mutations caused frameshifts in the open reading frame, predicted to produce truncated, non-functional proteins of 78/256 amino acids in Fgf3 and 144/202 amino acids in Fgf10a (Figure 1B).

**Figure 1.**
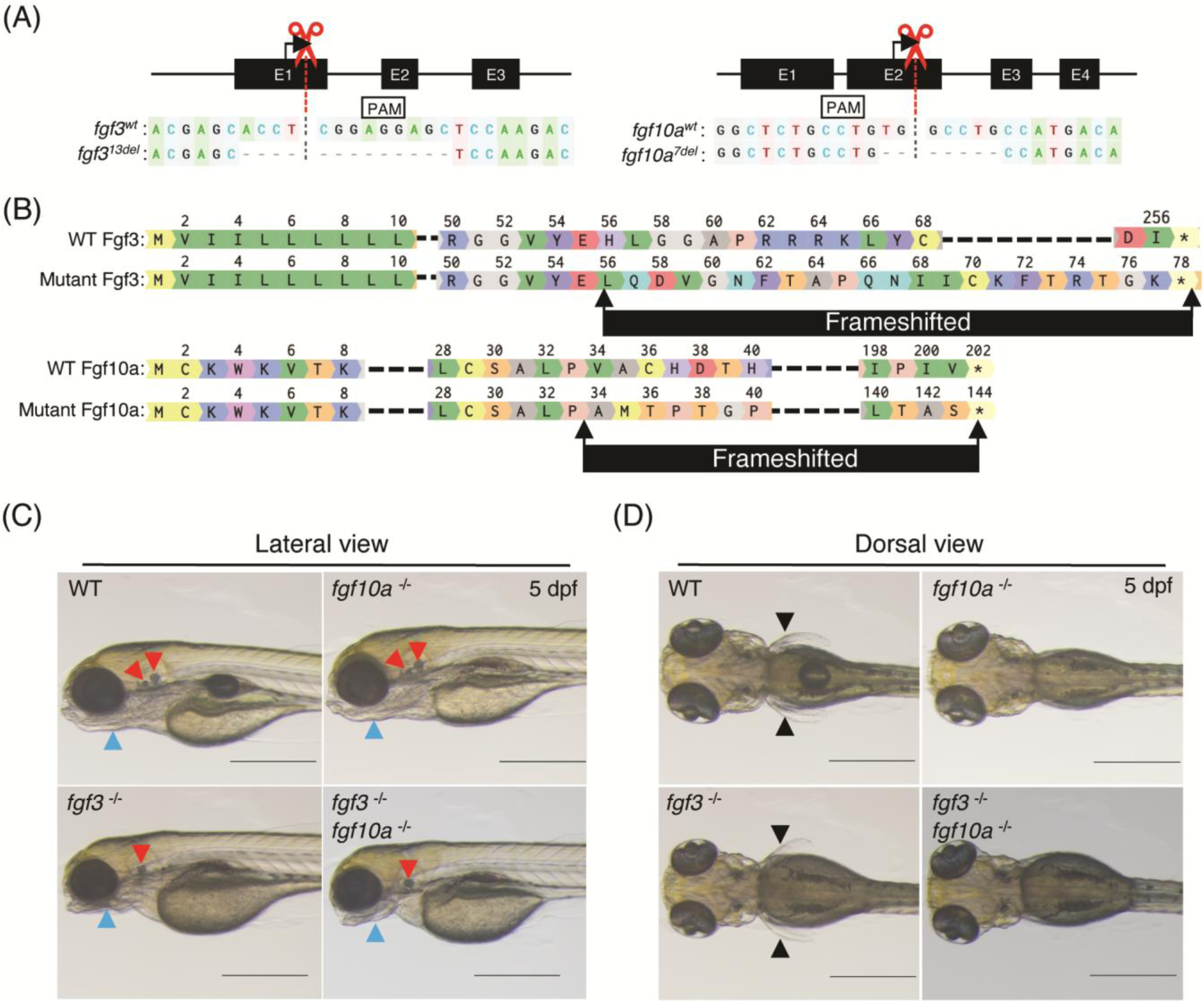
Knockout strategy and phenotypes of *fgf3*,*fgf10a* double mutant embryos. (A) Schematic representation of *fgf3* and f*gf10a* coding sequences, highlighting CRISPR-Cas9 cutting sites, protospacer adjacent motifs (PAM), and mutations in both genes. (B) Schematic diagrams of the Fgf3 and Fgf10a amino acid sequences in wildtype and mutant forms, showing frameshifts and truncated regions. (C-D) Brightfield microscope images of 5 dpf larva morphologies including (C) fused otolith and malformed jaw, and (D) absence of pectoral fins in wildtype, single and double *fgf3,fgf10a* mutant larva.

The loss of function of Fgf3 and Fgf10a was validated based on phenotypes previously described^17–20^. Loss of Fgf3 function caused otolith fusion within the otic vesicles and malformed jaws (Figure 1C), while loss of Fgf10a function resulted in absent pectoral fins (Figure 1D). The fgf3,10a double mutant exhibited both phenotypes, evident by 3 dpf, served as the basis for phenotypic classification before downstream experiments. In addition, Fgf3 and 10a were previously reported to work coordinately in regulating the migration of primordium during lateral line development^15,21^. We observed similar primordium migration delay phenotype in our in-house generated fgf3,10a double mutants (Supplementary Figure 1). Interestingly, extended observations showed that despite the delayed primordium migration, the primordium eventually reached the posterior end of the embryo, and the mutants were able to form a lateral line consisting of neuromasts and nerves (Supplementary Figure 1 A 72hpf, Figure 2B). Together, these results indicate that the *fgf3* and *fgf10a* mutants generated in this study phenocopy previously reported loss-of-function mutants, supporting that Fgf3 and/or Fgf10a function is abolished in these mutants.

**Figure 2.**
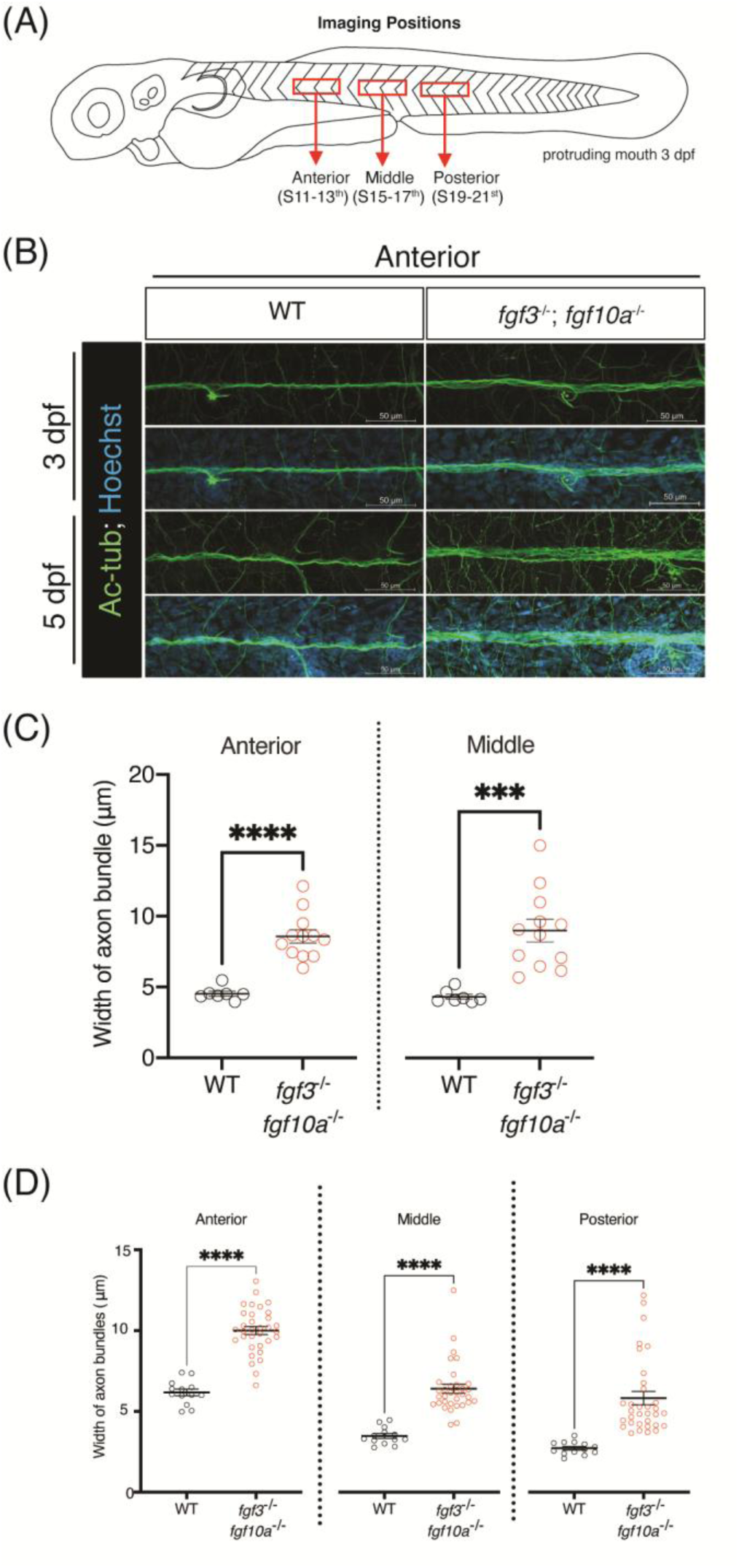
Confocal images of immunofluorescence staining visualizing axonal structures in wildtype and *fgf3*,*fgf10a* double mutants. (A) Illustration of imaging regions: anterior (somites 11-13^th^), middle (somites 15-17^th^) and posterior (somites 19-21^st^). (B) Confocal images of axons and cell nuclei labelled with anti-acetylated tubulin antibody and Hoechst, respectively in anterior regions of wildtype (WT) and *fgf3*,*fgf10a* double mutants at 3 and 5 dpf. Anterior is oriented to the left. Scale bars: 50 μm. (C-D) Mean width of axon bundles in anterior, middle and posterior regions in wildtype (WT) and *fgf3*,*fgf10a* double mutants at 3dpf (C) and 5 dpf (D). Each dot represents a single larva. Data are represented as mean ± S.E.M and were analysed using one-way ANOVA with post-hoc multiple comparisons, *** *p* < 0.005, **** *p* < 0.001.

### 2.2 *fgf3,10a* double mutants exhibit axonal defasciculation in the lateral line

Previous studies have reported that Fgf3 and Fgf10a are required for primordium migration and epithelialization during lateral line development^15,22^. However, it has remained unclear whether Fgf3 and Fgf10a are also required for the development of lateral line nerves, one of the major components of the lateral line system. To address this question, we performed whole mount immunostaining with anti-acetylated tubulin antibody to visualize the lateral line neurons in *fgf3, 10a* double mutants (Figure 2). We found that axon bundles in *fgf3*,*10a* double mutants appeared wider compared to those in wild type siblings (Figure 2A, B). We then performed quantitative analysis by measuring axon bundle width (the distance between the outermost axonal fibers) at 3 and 5 dpf. This analysis revealed statistically significant increases in axon bundle width in *fgf3*,*10a* double mutants at both 3 and 5 dpf across all positions examined (Figure 2C, D), as well as compared to other genotype siblings (Supplementary Figure 2), supporting our initial observation.

Next, we asked whether the increased axon bundle width might be due to an increase in neuronal cell number in *fgf3,10a* double mutants, given that *fgf3* mutants have been reported to induce hair cell proliferation in the zebrafish lateral line via Fgf and Notch signalling^23^. We therefore employed Tg[*HuC*: *kaede*] to visualize and quantify the number of cell body of lateral line neurons (called lateral line ganglion) at 2 and 3 dpf (Figure 3A, B). We found no statistically significant difference in the number of cell body in posterior lateral line ganglion between *fgf3,10a* double mutants and wild type embryos at both 3 and 5 dpf (Figure 3C). Taken together, these results suggest that the increased axon bundle width reflects loosening of axonal fasciculation rather than an increase in neuronal cell number.

**Figure 3.**
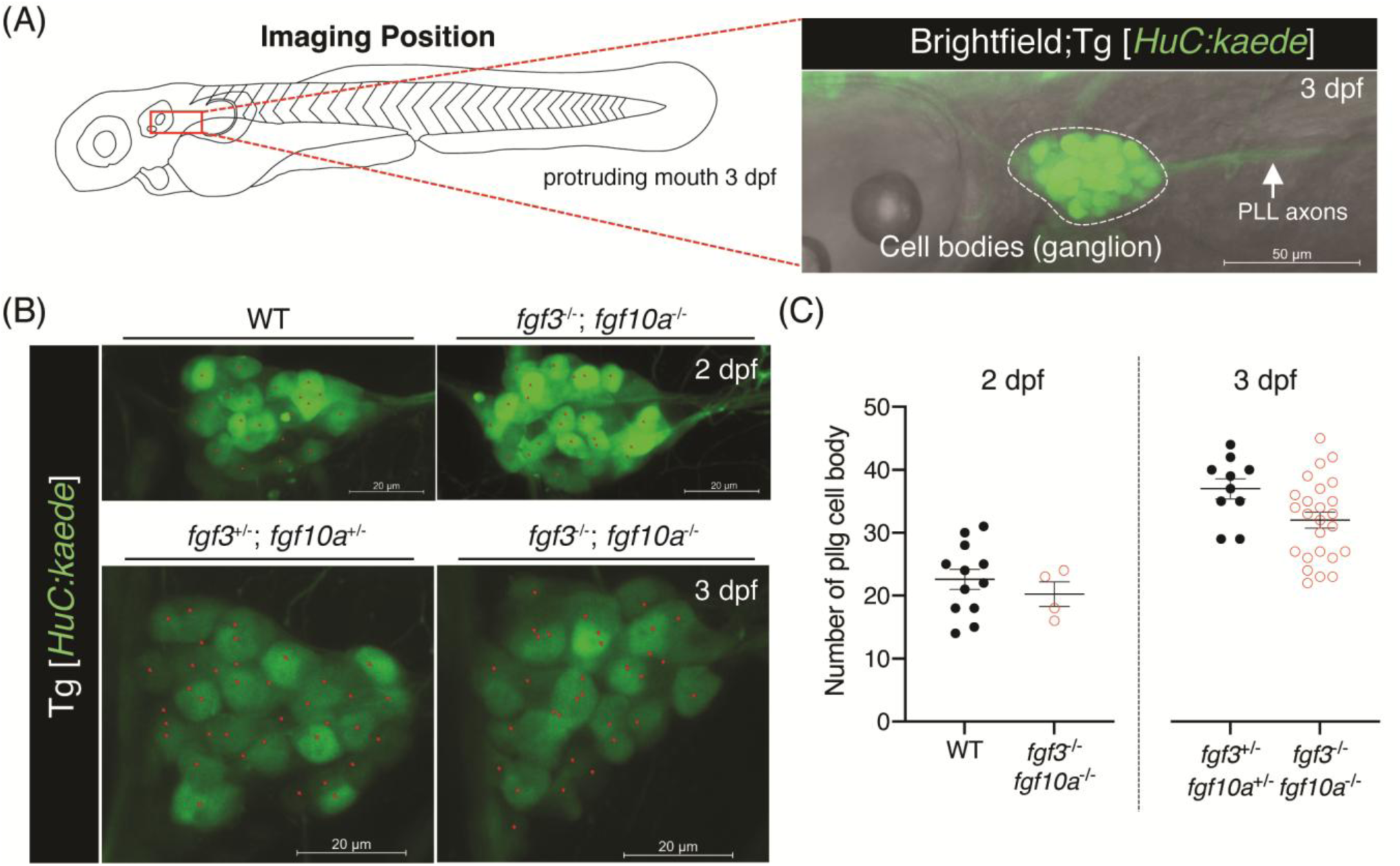
Visualization of posterior lateral line (pll) ganglion in wildtype (WT), heterozygous mutants and *fgf3*,*fgf10a* double mutants. (A) Schematic showing the imaging location of pll neuronal cell bodies. White dotted line outlines the ganglion, and white arrow indicates pll axons. Anterior is to the left. Scale bars: 50 μm. (B) Confocal images of pll ganglia in Tg[*HuC:kaede*] WT, heterozygous mutant and *fgf3*,*fgf10a* double mutants at 2 and 3 dpf. Scale bars: 20 μm. (C) Quantification of pll ganglion cell body in WT and *fgf3*,*fgf10a* double mutants at 2 dpf, and heterozygous mutants and *fgf3*,*fgf10* double mutants at 3 dpf. Each dot represents an individual larva. Data represent mean ± S.E.M.

### 2.3 Increased Schwann cell number in *fgf3,10a* double mutants

Schwann cells, a type of glial cell, are closely associated with peripheral axons. During early axon extension, Schwann cell precursors migrate along axons and differentiate into myelinating Schwann cells, which enwrap axons to maintain axon fasciculation^10,24^. Thus, we hypothesized that the axonal defects observed in *fgf3,10a* double mutants might be mediated by Schwann cells. To examine this possibility, we first analyzed the expression of Schwann cell markers, including *sox10* (sex determining region Y(SRY)-box transcription factor 10, expressed in mature Schwann cell and its precursor) and *mbpa* (myelin basic protein a, expressed in mature myelinating Schwann cells) using whole mount *in situ* hybridization (WISH). We found that *sox10* expression in control siblings was detected along the horizontal myoseptum at 3 dpf, corresponding to the pll position (Figure 4). In contrast, *fgf3,10a* double mutants exhibited a dorsoventrally expanded *sox10* expression domain along the lateral line (Figure 4), suggesting an increase in the number of *sox10*-expressing Schwann cells in the mutants. Meanwhile, the width of the *mbpa* expression domain in *fgf3,10a* double mutants was comparable to that in control siblings. This observation suggests that the expanded *sox10* expression domain reflects an increase in Schwann cell precursor populations rather than mature myelinating Schwann cells.

**Figure 4.**
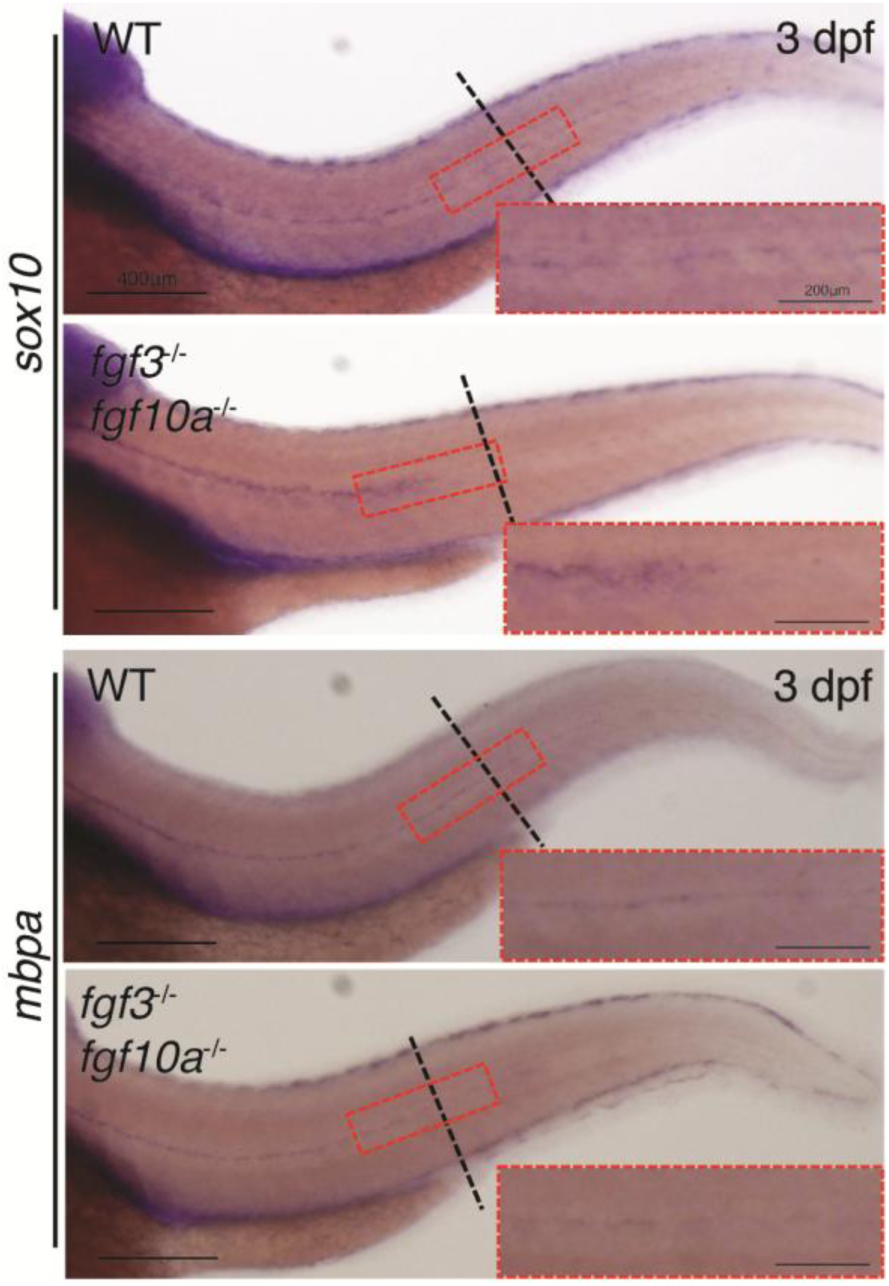
Expression of *sox10* and *mbpa* in zebrafish posterior lateral line in wildtype and *fgf3,fgf10a* double mutants at 3 dpf. Dotted lines represent the location of the 16th somite, and images within red dotted rectangle are magnified views of the indicated regions. Anterior is to the left. Scale bars = 400 and 200 μm as indicated.

To further characterize the increase in Sox10-positive (So×10^+^) cells observed in *fgf3,10a* double mutants, we visualized the Schwann cells by whole-mount immunostaining using anti-Sox10 antibody. At 5 dpf, we found that *fgf3,10a* double mutants exhibited significantly higher number of So×10^+^ cells, with nearly an 18-34% increase in So×10^+^ Schwann cell numbers compared to wild type and other genotype siblings (Figure 5). These results suggest that Fgf3 and Fgf10a are required to maintain the proper number of Schwann cell precursors in the lateral line. Importantly, we found no morphological defects in the spinal cord of *fgf3,10a* double mutants, further confirming that this defect is specific to the pll (Supplementary Figure 3)

**Figure 5.**
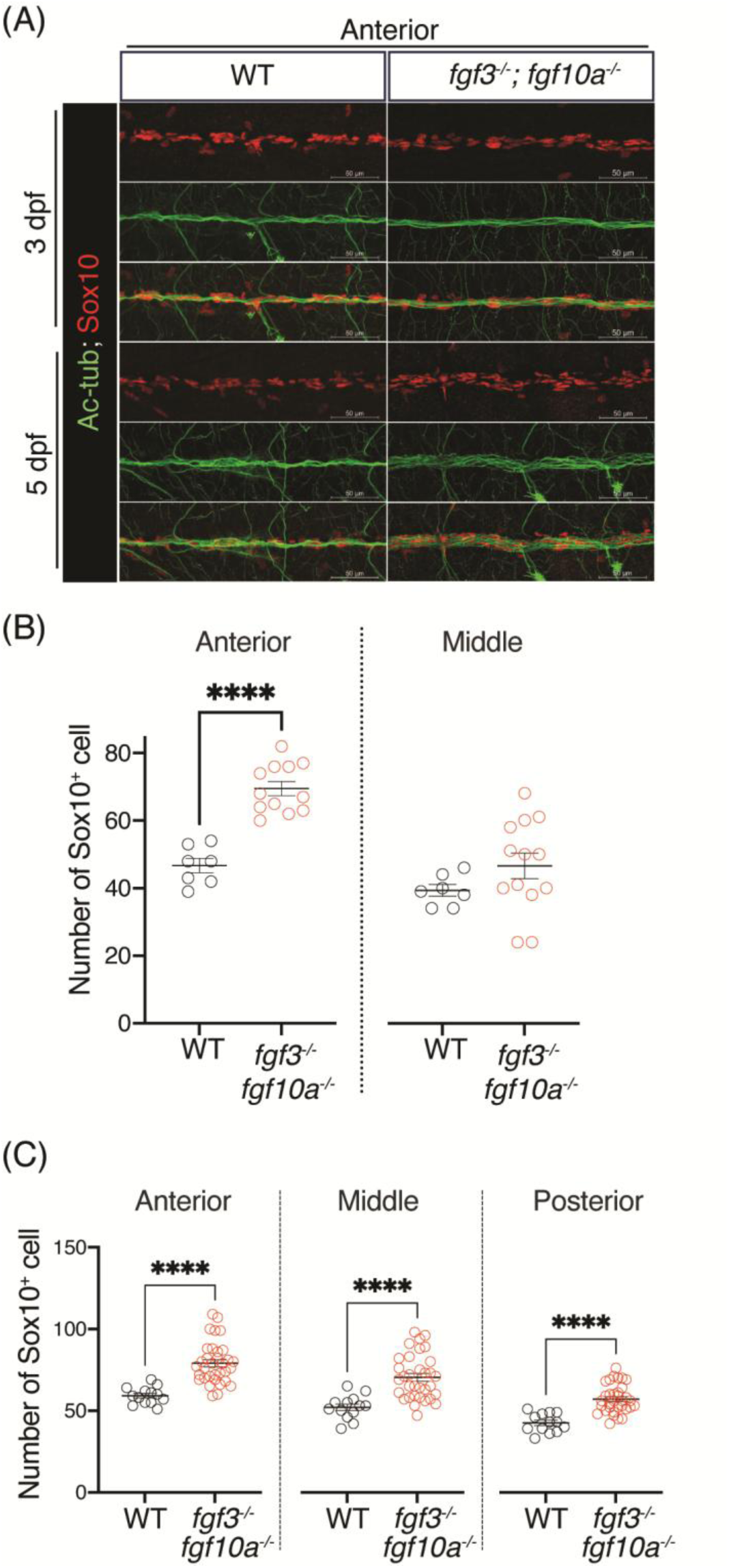
Confocal images of immunofluorescence staining visualizing Schwann cells and axons in wildtype and *fgf3*;*fgf10a* double mutants. (A) Confocal images of So×10^+^ cell and axons labelled with anti-Sox10 and anti-acetylated tubulin antibodies at 3 and 5 dpf in anterior region in wildtype (WT) and *fgf3*,*fgf10a* double mutants. Anterior is to the left. Scale bars: 50 μm. (B-C) Quantification of the mean number of So×10^+^ cell in anterior, middle, and posterior regions in wildtype (WT) and *fgf3*,*fgf10a* double mutants at 3 dpf (B) and 5 dpf (C). Each dot represents a single larva. Data represent mean ± S.E.M and were analysed by one-way ANOVA with Tukey’s post-hoc multiple comparisons, **** *p* < 0.001.

At 5 dpf, Sox10⁺ cell numbers were significantly increased across all regions of pll (anterior, middle, and posterior) in *fgf3,10a* double mutants compared to wild type siblings. In contrast, at 3 dpf, the number of Sox10⁺ Schwann cells was significantly increased only in the anterior region, whereas in the middle region, Sox10⁺ cell numbers tended to be higher in the mutants, but the difference was not statistically significant (Figure 5). At earlier developmental stages, no clear difference in the number of Sox10⁺ cells associated with the lateral line was observed between genotypes (31 hpf; Supplementary Figure 4). These results suggest that the increase in Sox10⁺ cells progress over developmental time and, in the middle region, becomes evident after 3 dpf. To capture the onset of this increase, we focused on the middle region just before and around 3 dpf for subsequent live imaging analysis.

### 2.4 Live imaging reveals higher frequency of glial cell proliferation and movements into the interaxonal spaces in *fgf3,10a* double mutants

Given the increase in Schwann cell number observed, we further examined how changes in Schwann cell number affect axonal organization *in vivo* in *fgf3,10a* double mutans. To assess this, we cloned zebrafish 7k bp upstream enhancer of *sox10* to generate glial cell reporter line Tg[*sox10:TagRFP*] in *fgf3,10a* heterozygous mutants. Together with Tg[*HuC:Kaede*] we visualized the interaction of lateral line axons and Schwann cell with *in vivo* live-imaging at 68–75 hpf in the middle region of the lateral line (Figure 6A), corresponding to the period just before and around 3 dpf, when Sox10⁺ cell numbers start to show an increasing trend in this region. During the imaging period, So×10^+^ cell in *fgf3,10a* double mutants divided approximately five times more frequently (5.3 ± 0.3 cells/3h) than in wild type (1.0 cell/3h) at the proximal region of lateral line neuron (Figure 6B,C, Supplementary Movie 1, 2).

**Figure 6.**
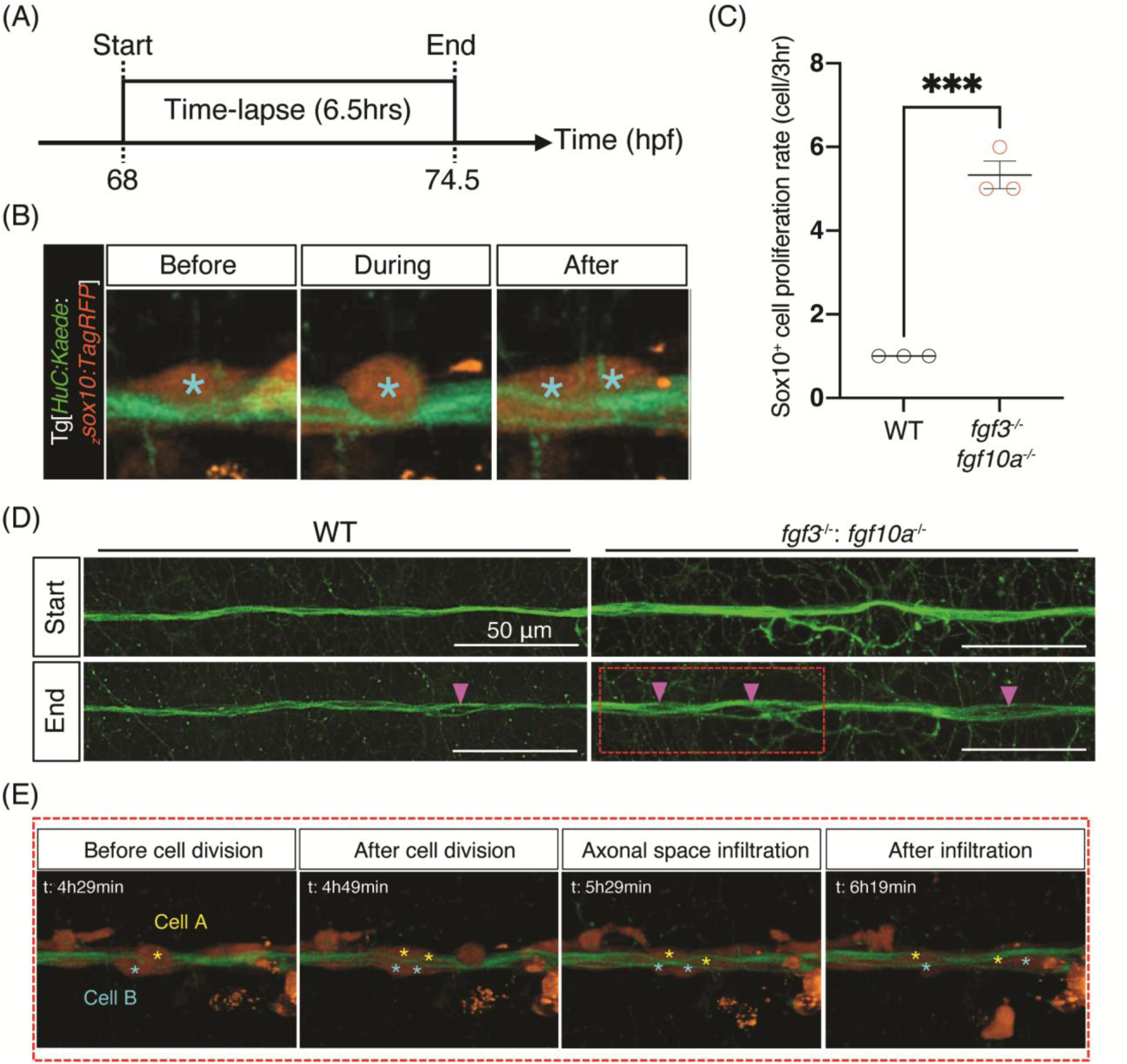
Timelapse *in vivo* imaging of Schwann cell proliferation and axon defasciculation at 3 dpf. (A) Schematic diagram depicting the time-lapse *in vivo* observation period in wildtype (WT) and *fgf3,fgf10a* double mutants. (B) Snapshot frames showing Schwann cell division (before, during and after cell division) obtained from time-lapse imaging video. (C) Quantification of Schwann cell division frequency in WT and *fgf3,fgf10a* double mutants. Each dot represents a single video. Data represent mean ± S.E.M. Statistical significance was assessed using Student’s t-test, with *** *p* < 0.005. (D) Frames from time-lapse *in vivo* imaging at the start and end of observation period in WT and *fgf3,fgf10a* double mutants. Anterior is to the left. Scale bars: 50 μm. Magenta arrowheads indicate the site of axon defasciculation. (E) Magnified snapshot frames (red dotted box) of time-lapse video showing Schwann cell infiltration (Cell A and B marked by yellow and blue asterisks) following division, leading to axonal defasciculation in *fgf3,fgf10a* double mutants.

Live imaging further revealed that newly divided So×10^+^ daughter cells migrated along axons and subsequently infiltrated interaxonal spaces (Figure 6E; Supplementary Movie 2, yellow arrowhead). This infiltration was accompanied by expansion of interaxonal spacing and disruption of axonal fasciculation (Figure 6D, arrowheads). Although such events were occasionally observed in wild type larvae, they occurred more frequently in *fgf3*,*10a* double mutants (Supplementary Movie 2), consistent with the increased proliferation rate of Schwann cells in these mutants. These results demonstrate that excessive Schwann cell proliferation drives interaxonal infiltration of daughter cells, thereby disrupting axonal bundling in *fgf3,10a* double mutants.

### 2.5 Nrg1/ErbB signaling involves in regulating Schwann cell overproliferation in *fgf3,10a*

### double mutants

Neuregulin 1 (Nrg1) is a member of the epidermal growth factor ligand family secreted by axons and regulates Schwann cell proliferation through binding to ErbB2/3 receptors^25–27^. Nrg1 is also known to play roles in regulating Schwann cell migration^28,29^, proliferation^24^, and myelination^29,30^ in zebrafish pll. Therefore, we hypothesized that Nrg1 expression might be altered in *fgf3*,*10a* double mutants. We first visualized *nrg1* expression in the pll ganglion using whole-mount *in situ* hybridization (WISH), and observed upregulation of *nrg1* in *fgf3,10a* double mutants compared to wild type at 3 dpf (posterior to the otic vesicle [OV]; magnified region indicated by the white dotted box) (Figure 7).

**Figure 7.**
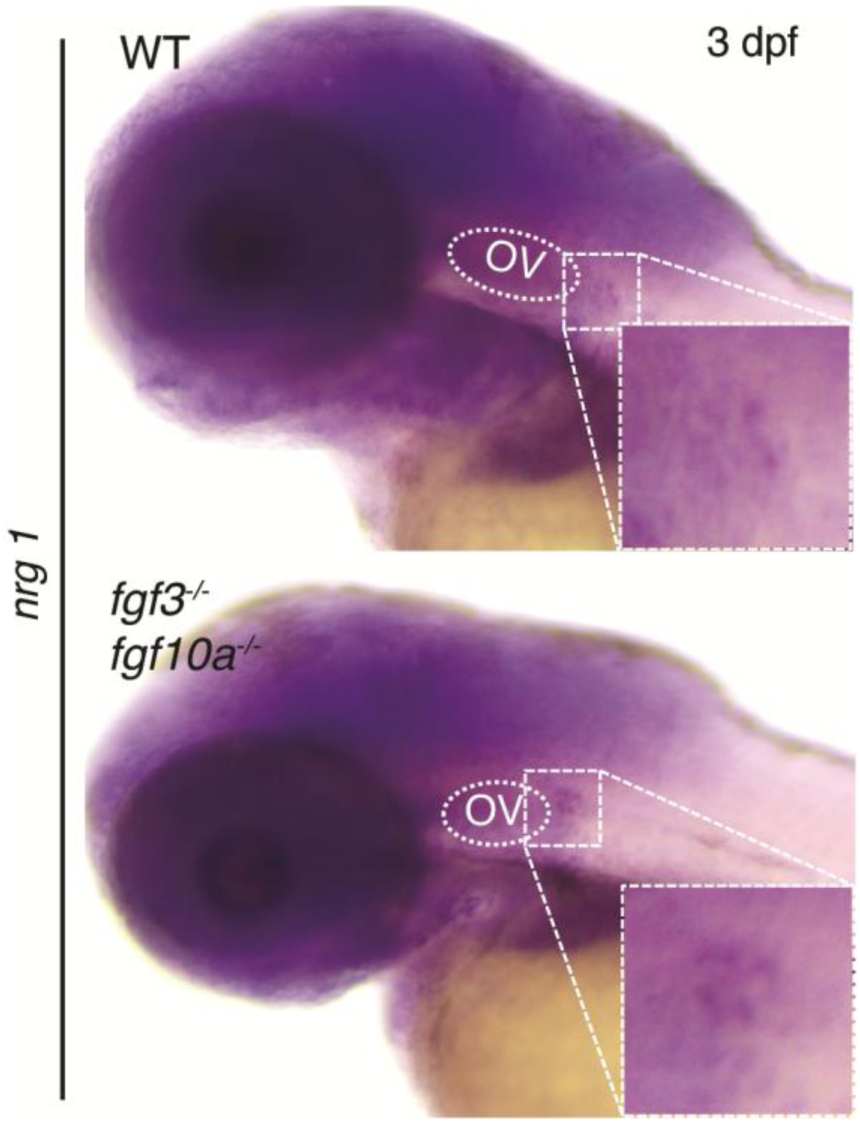
*nrg1* expression in wildtype (WT) and *fgf3,fgf10a* double mutants at 3 dpf. The small dotted box indicates the location of pll ganglion. Images within the dotted line rectangle show a magnified view of the pll ganglion area; OV, is otic vesicle.

To examine whether Nrg1/ErbB signaling contributes to the increased proliferation of Sox10⁺ cells in *fgf3,10a* double mutants, we applied the ErbB receptor antagonist AG1478 to inhibit Nrg1/ErbB signalling and determine whether limiting Nrg/ErbB signalling can rescue Sox10⁺ cell proliferation in *fgf3,10a* double mutants. AG1478 treatment from 2-5 dpf reduced the number of So×10^+^ cells by 56% in *fgf3,10a* double mutants (42.8 ± 2.7 cells) compared to DMSO-treated group (97.7 ± 4.6 cells) (Figure 8, Supplementary Figure 5). Meanwhile, the mean width of the axon bundle was also significantly reduced in AG1478-treated group (6.5 ± 0.8 μm) compared to the DMSO-treated group (9.5 ± 0.7 μm), indicating partial rescue of the axonal defasciculation phenotype (Figure 8, Supplementary Figure 5). These results suggest that limiting the Nrg1/ErbB signalling effectively restricts So×10^+^ cell proliferation and ameliorates axonal defasciculation in *fgf3,10a* double mutants.

**Figure 8.**
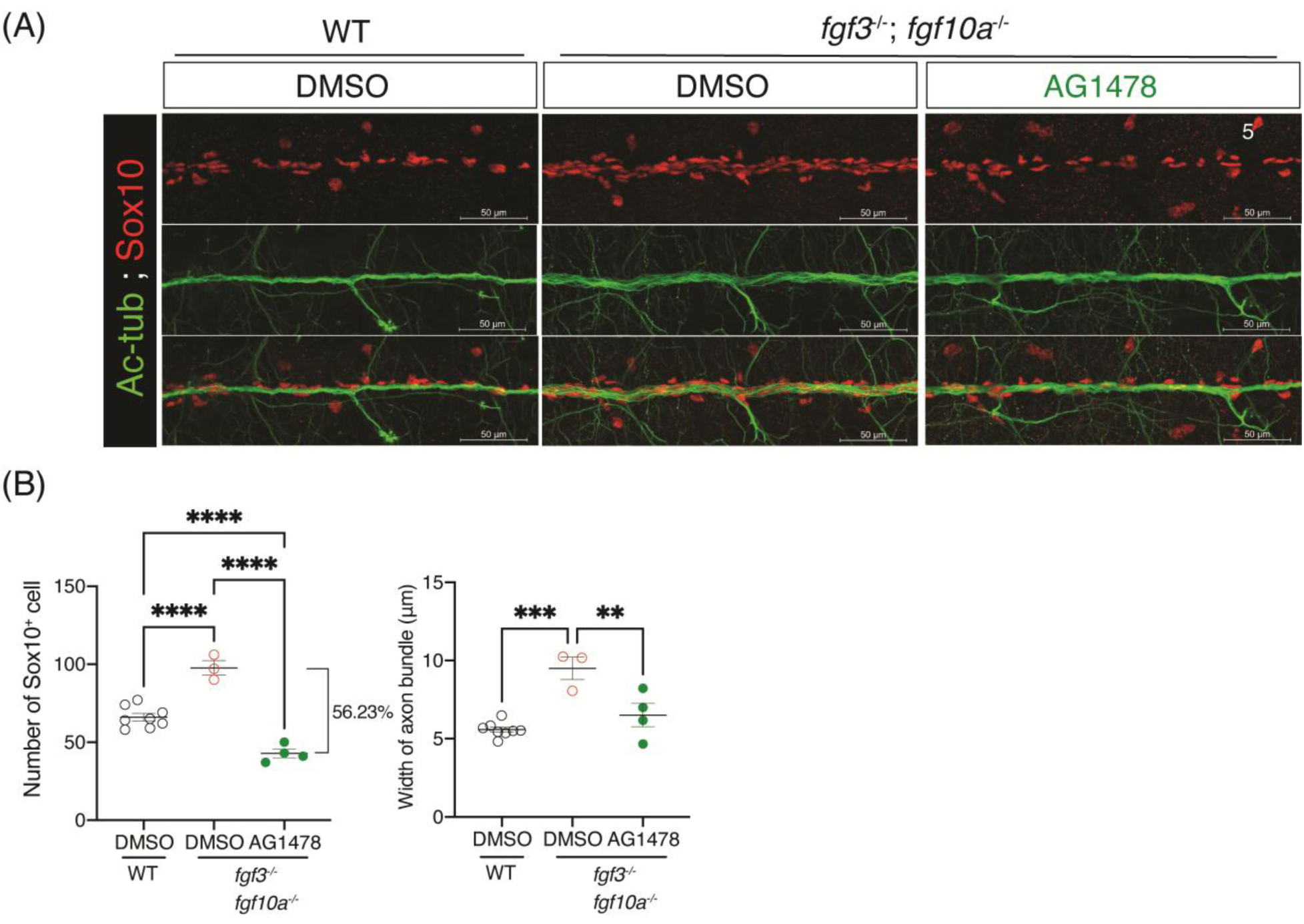
Confocal imaging of immunofluorescence staining visualizing Schwann cell and axon at 5 dpf following 2-3 dpf or 2-5 dpf AG1478 treatments. (A) Confocal images of So×10^+^ cells and axons labelled with anti-Sox10 anti-acetylated tubulin antibodies at 5 dpf in the anterior region of 2-5 dpf DMSO treated wildtype (WT), DMSO treated *fgf3*,*fgf10a* double mutants, and AG1478 treated *fgf3,fgf10a* double mutants. Anterior is to the left. Scale bars: 50 μm. (B) Quantification of the mean number of So×10^+^ cell and mean axon bundle width at 5 dpf in the anterior region of DMSO treated wildtype (WT), DMSO treated *fgf3,fgf10a* double mutants, and AG1478 treated *fgf3,fgf10a* double mutants for 2-5 dpf treatment duration. Each dot represents a single larva. Data represent mean ± S.E.M. One-way ANOVA with post-hoc multiple comparisons was used to assess statistical significance, \*\**p* < 0.01, *** *p* < 0.005, **** *p* < 0.001.

To determine whether the upregulation of *nrg1* in pll ganglion influences Schwann cell number, *nrg1-typeIII* was overexpressed specifically in neuronal cells using the pan-neuronal *HuC* promoter. Enhanced green fluorescence protein (EGFP) served as a reporter for Nrg1 overexpression (Figure 9). Larvae expressing EGFP in neurons were classified as Nrg1^+^, while their non-EGFP siblings served as controls. Schwann cell and axonal morphology were visualized using immunofluorescence staining with anti-Sox10 and anti-acetylated tubulin antibodies, as previously described. Results showed a significantly increase by 56% in So×10^+^ cell number (98.2 ± 10.1 cell) and 34% in axon bundle width (7.5 ± 0.9 μm) in Nrg^+^ larvae compared to controls (62.9 ± 7.0 cell, and 5.6 ± 0.3 μm, respectively) (Figure 9C). This supports our hypothesis that the Schwann cell and axonal phenotypes observed in *fgf3,10a* double mutants resulted from *nrg1* upregulation.

**Figure 9.**
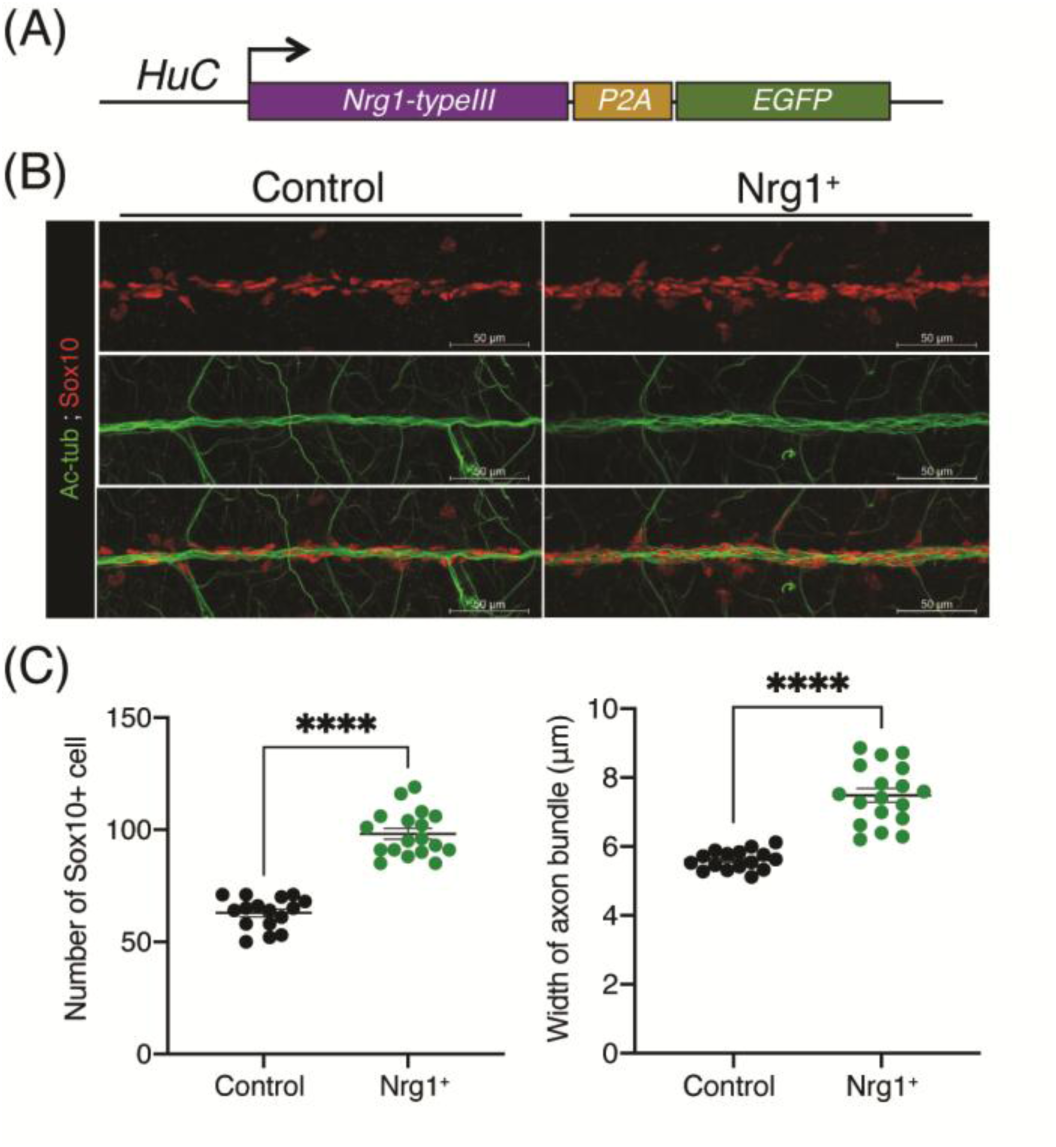
Neuronal cell-specific overexpression of Nrg1-typeIII and its effects on Schwann cell and axonal morphology. (A) Schematic of the DNA construct used to generate the Nrg1-typeIII overexpressing line Tg[*HuC:nrg1-typeIII-P2A-EGFP*]. (B) Confocal images showing So×10^+^ cell and axons labelled with anti-Sox10 and anti-acetylated tubulin antibodies at 5 dpf in anterior region of EGFP positive (Nrg^+^) and control larva. Anterior is oriented to the left. Scale bars: 50 μm. (C) Quantification of the mean number of So×10^+^ cell and mean width of axon bundle in anterior region. Each dot represents a single larva. Data represent mean ± S.E.M and were analysed by student t-test, \*\*\*\**p* < 0.001.

## 3 DISCUSSION

### 3.1 Fgf3 and Fgf10a maintain axonal fasciculation through the suppression of excessive Schwann cell proliferation and infiltration

In this study, we analysed *fgf3,10a* double mutants, which have been previously reported to show defects in primordium migration and differentiation. In addition to the previously described delay in primordium migration, we identified a novel phenotype in which pll axons running along the horizontal myoseptum exhibited clear defasciculation.

Previous studies have shown that Schwann cells and their glial precursors are essential for the formation and maintenance of axonal fasciculation in the lateral line nerve^10^. In those studies, the absence of Schwann cells resulted in defective axonal bundling. In contrast, our analysis revealed the opposite situation in *fgf3,10a* double mutants, where the number of So×10^+^ Schwann precursor cells was increased compared with wild type embryos. Moreover, regions in which Sox10-positive cells were located between axons corresponded to areas where axon bundles appeared expanded and loosely organized.

Live imaging further revealed enhanced Schwann cell proliferation in the mutants. Notably, daughter cells produced by the division of So×10^+^ cells along axons frequently migrated into the spaces between neighboring axons, a behavior we define here as “infiltration.” This infiltration increases the spacing between adjacent axons, leading to loosening of axonal bundles. These results suggest that Fgf3 and Fgf10a maintain axonal fasciculation during pll development by restricting Schwann cell proliferation and preventing excessive infiltration into axon bundles.

During normal peripheral nerve development, Schwann cell precursors proliferate while migrating along axons. They subsequently become immature Schwann cells that enter axon bundles and separate individual axons through the process known as radial sorting, after which they move to the axonal surface and differentiate into myelinating Schwann cells. In the present study, we found that proliferation of So×10^+^ cells persists even after their migration along the lateral line nerve has been completed in *fgf3,10a* double mutants. Consequently, excess Schwann cells infiltrate into axon bundles, which likely disrupts the radial sorting process and leads to defective axonal fasciculation.

### 3.2 Nrg1/ErbB signaling contributes to Schwann cell overproliferation in *fgf3,10a* double mutants

Members of the EGF ligand family, particularly Neuregulin 1 (Nrg1), play important roles in peripheral nerve development by regulating Schwann cell behaviors through the ErbB2/3 receptor complex ^27,31–34^. In zebrafish, Nrg1 type-III is expressed in lateral line axons and regulates Schwann cell migration, proliferation, and myelination^24,28–30^. In the present study, *in situ* hybridization revealed a modest increase in *nrg1* expression in the pll ganglion of *fgf3,10a* double mutants, suggesting a potential involvement of Nrg1/ErbB signaling.

Consistent with this possibility, pharmacological inhibition of ErbB signalling using AG1478 reduced the elevated number of Sox10⁺ cells and partially rescued the axonal defasciculation phenotype in *fgf3,10a* double mutants. In addition, neuronal overexpression of Nrg1-typeIII increased Sox10⁺ cell number and axon bundle width. Together, these observations suggest that enhanced Nrg1 signaling promotes Schwann cell proliferation and may contribute to the disruption of axon fasciculation.

*fgf3* and *fgf10*a are expressed in the pre-migratory and migrating lateral line primordium as well as in craniofacial tissues surrounding the pll ganglion^14,18^. Because these ligands are secreted factors, lateral line neurons and associated Schwann cells are positioned in close proximity to potential Fgf sources during early development. In the present study, we observed an modest upregulation of *nrg1* expression in the pll ganglion of *fgf3*,*10a* double mutants, raising the possibility that Fgf signaling may modulate Nrg1 signaling in this region. However, the direct cellular targets of Fgf3 and Fgf10a in this context remain unclear. It also remains to be determined whether Fgf3 and 10a act directly on lateral line neurons, Schwann cells, or surrounding tissues to influence Nrg signaling.

In conclusion, we revealed novel roles of fibroblast growth factors in regulating sensory nerve organization and Schwann cell development in the zebrafish lateral line system. Our results suggest that Fgf3 and Fgf10a are required to ensure proper Schwann cell differentiation and proliferation, thereby maintaining normal axonal fasciculation. Loss of Fgf3 and Fgf10a leads to overproliferation of Schwann cells in the pll and promotes axonal defasciculation, likely through infiltration of daughter Schwann cells into interaxonal spaces. Our data also suggest that Nrg1/ErbB signaling may contribute to this process. While additional studies are needed to clarify how Fgfs regulate axon-derived Nrg1 expression in sensory nervous system, our findings highlight the importance of Fgf signaling in regulating Schwann cell development and axonal organization during pll development.

## 4 EXPERIMENTAL PROCEDURE

### 4.1 Animals

Zebrafish (*Danio rerio*) used in this study were maintained in NAIST zebrafish facility under standard condition at 28.5°C with a 14-hour light/ 10-hour dark cycle. Zebrafish were bred according to the method described in the zebrafish book^35^ and eggs collected were kept in embryo medium (RO water with 0.03% artificial salt and 0.00003% methylene blue) at 28.5°C until desired developmental stage. Embryos were staged according to zebrafish developmental stages as previously described^36^. All zebrafish experiments were approved by Nara Institute of Science and Technology’s Animal Studies Committee

### 4.2 CRISPR/Cas9 design and crRNA synthesis

Mutations were introduced into the first coding exon of *fgf3* and *fgf10a* genes using the CRISPR/Cas9 system as previously described in zebrafish^37^. Briefly, gRNAs targeting the first exon of both genes were designeduding CrisprDirect and Crisprscan^38–40^. We synthesized gRNAs using primers composed of a T7 promoter (**Supplementary Table 1**) according to GeneWeld protocol^41^ using MEGAscript T7 Transcription kit (Ambion, USA). The gRNA was then purified with miRNeasy Qiagen kit (Qiagen, USA) according to manufacturer’s protocol and the concentration was measured with NanoDrop One/OneC Spectrophotometer (ThermoFischer Scientific, USA). gRNA was stored at −80°C until further use.

### 4.3 Generation and genotyping of Knock-out Zebrafish

1-10 nl of the injection mixture containing 25 ng/ml of gRNA, 100 ng/ml of *Cas9* mRNA, and 0.05% of phenol red diluted in Milli-Q water was injected into one-cell stage wild type zebrafish embryos using FemtoJet microinjector (Eppendorf, Germany) equipped with dissecting microscope (Olympus, Japan). Healthy injected larva (referred to here as crispans) were raised in the system until adulthood for founder selection.

Crispans were out crossed with wild type fish, and the embryos were used for founder selection. Ten embryos were pooled in a single PCR tube for genomic DNA extraction. Approximately 300-400 bp regions containing the gRNA target sites were amplified by PCR using DreamTaq Green PCR Master Mix (2X) (Thermo Fisher Scientific™, USA) with primers (zFgf3_Fw1_2 and zFgf3_Rv for *fgf3*, and FGF10a_R2_Fw and FGF10a_R2_Rv for *fgf10a*) in a total volume of 10 μl. Primer sequences are listed in **Supplementary Table 1**. The amplicons were sent for sanger sequencing by GeneWiz Inc, Japan to detect mutations. The efficiency and type of CRISPR/Cas9 induced mutation were evaluated using ICE analysis (https://ice.synthego.com/#/). The crispan embryos containing the desired mutations were used to generate the next generation of germline mutant zebrafish (F1). F1 larvae were raised to adulthood, and mutations were verified again by fin clip genomic DNA extraction and sanger sequencing.

For further verification, the amplicons of the mutant genes were cloned into a vector using pGEM®-T Easy Vector kit (Promega, USA) following the manufacturer’s protocol and transformed into Competent *E. coli* DH5α (Nippon gene, Japan) with Blue/White selection using X-Gal (50 μl/plate). The plasmid containing mutant DNA was verified with sanger sequencing and used for PCR genotyping primer design. All PCR genotyping primers were designed using Primer3Plus software (https://www.primer3plus.com/index.html)^42^ spanning 300-400 bp of the mutation site with annealing temperatures of 66.5°C for *fgf3*^wt/13del^, and 65°C for *fgf10a*^wt/7del^, primer pairs are shown in **Supplementary Table 2**.

### 4.4 Genomic DNA extraction and PCR genotyping

The embryos were raised to adulthood, and the caudal fin (3-5 mm) were excised with 70% ethanol disinfected scissors. Clipped fin or dechorionated embryo were put into PCR tube and genomic DNA was extracted using 40 μl Fin Clip buffer (10 mM Tris-HCl (pH8.4)), 50 nM KCl, 1.5 mM MgCl_2_, 0.3% Tween, 0.3% NP-40). The samples were incubated at 95°C for 20 min, then added 4 μl of 10 mg/ml proteinase K (Nacalai Tesque, Japan) and incubated for 2 hours at 55°C followed by 95°C for 20 min to denatured proteinase K. The DNA samples were kept at −30°C until further use.

### 4.5 Construction of plasmids for zebrafish transgenic line

DNA fragment containing *sox10* enhancer region (6955 bp) was amplified from zebrafish genomic DNA using the forward primer: LR_Apa1_sox10_prom_Fw and the reverse primer: TagRFP_Koz_Agel_zSox10_prom_Rv, which were designed according to enhancer region as predicted^43^. The TagRFP sequence (746 bp) was amplified using the forward primer: TagRFP_Fw and the reverse primer: Sv40_Cla1_Ascl_TagRFP_Rv. All amplicons were undergone restriction enzyme digestion and ligated to *pT2aL200R150G* backbone vector using Ligation high Ver.2 (Toyobo) following manufacturer’s protocol, resulting *pT2a-_z_sox10:TagRFP* plasmid.

The DNA fragment containing *nrg1-typeIII* coding sequence with a flanking homologous region (2146 bp) was amplified from zebrafish cDNA using the forward primer: Gib_Agel_Koz_nrg1_typeIII_Fw and the reverse primer: Gib_P2A_nrg1_typeIII_Rv. The EGFP sequence was amplified with a flanking homologous region (802 bp) using the forward primer: Gib_P2A_EGFP_Fw and the reverse primer: Gib_Sv40_Asc1_EGFP_Rv. The amplicons were then cloned to *pT2a-HuC:MCS* our inhouse generated backbone vector with Gibson assembly using NEBuilder® HiFi DNA Assembly Master Mix (NEB, USA) following the manufacturer’s protocol. All primers used are listed in **Supplementary Table 1**.

### 4.6 Generation of transgenic zebrafish line by Tol2 transposon system

The transgenic zebrafish lines in this study were generated by incorporating the targeted sequence into the wild type zebrafish genome using the Tol2 transposon system according to the method previously described^44^. The injection mixture containing plasmid (50 ng/μl) and transposase mRNA (50 ng/μl) in 0.2% phenol red was injected into one-cell stage wild type zebrafish embryos using a FemtoJet microinjector (Eppendorf, Germany) equipped with a dissecting microscope (Olympus, Japan). Each embryo received approximately 1-10 nl of injection mixture.

Embryos with integrated plasmid DNA were identified by screening embryos that express the fluorescence reporter protein using a DP70 light microscope (Olympus, Japan) equipped with a mercury lamp and GFP or RFP filter. Embryos with strong, positive signals in the correct cell types were selected and raised to adulthood. Injected founders (F0) carrying germline cells with the fluorescence reporter protein were identified by mating them with adult wild type zebrafish. The resulting embryos with positive signals were raised for the generation of a stable transgenic line.

### 4.7 Sample collection and fixation

Embryos were fixed with 4% paraformaldehyde/0.1% Triton/PBS overnight at 4°C, followed by dehydration with 100% methanol and stored in methanol at −30°C until the use for whole mount immunostaining or *in situ* hybridization.

### 4.8 Probe synthesis and whole-mount *in situ* hybridization (WISH)

A 6 dpf zebrafish cDNA pool was used to amplify fragments of targeted genes *via* PCR using KOD FX Neo (Toyobo, Japan) according to the manufacturer’s protocol. The primers designed to amplify the target genes are listed in **Supplementary Table 1**. Each forward primer contained a T3 promoter and reverse primer contained a T7 promoter to enable the synthesis of antisense or sense transcripts. Digoxigenin (DIG)-labelled RNA probes were synthesized from 200 ng of amplicon using DIG labelling kit (Roche) following the manufacturer’s protocol. The probes were denatured with 10 μl proteinase K (10 mg/ml) at 55°C for 30 minutes and precipitated with 100% ethanol, followed by washing with 75% ethanol and dissolved in hybridization buffer (60% formaldehyde, 5X SSC, 20% Tween 20 (Nacalai Tesque, Japan) and 50 mg/ml heparin (Sigma, USA). the quality and concentration of the probe were assessed by loading 0.5 μl RNA product onto a 1.5% agarose gel. All probed were kept at −30°C until further use.

Whole mount in situ hybridization was performed on fixed embryos as described elsewhere^45^ with modification as follows. Re-hydrated embryos were treated with 10 μg/ml proteinase K (Roche, Germany) for 5 minutes to enhance the probe and buffer penetration prior to pre-hybridization. The hybridization was then replaced with 300 ng/ml of in situ hybridization probe and incubated at 70°C overnight. The embryos were blocked with 10% sheep serum in TBST for 1 hour at room temperature followed by incubating in freshly prepared 1/2000 diluted anti-DIG alkaline phosphate (Roche, Germany) in 10% sheep serum/TBST overnight at 4°C. The mRNA expression was visualized with single color solution (22.5 μl NBT (Roche, Germany), 3.5 μl BCIP (Roche, Germany) in 1 ml NTMT solution) and incubated under dark condition at room temperature. After the colour developed, the samples were washed with PBT for 2-3 times to stop the reaction and kept in 4% PFA/PBST for long-term storage or examination with a light microscope.

### 4.9 Whole Mount Immunostaining

Dehydrated zebrafish embryos were treated with 3% H2O2 in methanol for 1 hour and rehydrated by sequential washes with 50%, 25% methanol/PBS, followed by PBS at 5-minute intervals, PBST twice at 10-minute intervals, and PBT three times at 5-minute intervals. To improve the permeability, the embryos were then washed with ultrapure water twice and immersed in pre-chilled 100% acetone for 8 minutes at −30°C. The embryos were then rinsed with ultrapure water twice to remove acetone and washed with PBT 3 times at 5-minute intervals. To inactivate endogenous peroxidases and increase the permeability of the cells, the embryos were incubated with 3% Triton X/1% DMSO in PBS at room temperature on a shaker for 1 hour. The embryos were then washed with MABDT three times at 5-minute interval at room temperature and blocked with 2% FBS/MABDT for three hours at room temperature on a shaker. The blocking buffer was replaced with a diluted primary antibody in 2% FBS/MABDT and incubated at 4°C on a shaker for 2 days. The concentration of antibody used was as follows: 1/300 anti-Sox10 (GeneTex, USA) and 1/750 anti-acetylated tubulin (Sigma-Aldrich, USA). After this, the embryos were washed with MABDT six times at 15-minute intervals at room temperature and incubated with a 1/1000 dilution of secondary antibody (Alexa Fluor 488, 594, 647) in 2% FBS/MABDT at 4°C on a shaker for 2 days. The samples were then washed with MABDT six times at 15-minute intervals at room temperature and incubated with a 1/1000 dilution of Hoechst in 4% PFA/PBS at 4°C overnight. Immuno-stained embryos were then washed with MABDT six times at 15-minute intervals at room temperature, followed by PBT three times at room temperature at 5-minute intervals, and then stored in 4% PFA/PBS at 4°C until confocal imaging.

### 4.10 Pharmacological treatment

The chorion of the embryos was removed by forceps at 24 hpf and subjected to pharmacological drug treatment for the desired duration. All inhibitors were diluted with dimethyl sulfoxide (DMSO) as stocks and used at the given concentration: 4 μM AG1478 diluted in embryo medium with 1X PTU. An equivalent volume of DMSO was used as a negative control for each treatment. All treatments were performed in a flat-bottom 24-well plate in the incubator at 28.5°C under dark conditions and the medium was replaced every 24 hours to maintain treatment efficiency.

### 4.11 *In vivo* time-lapse, confocal microscopy and light microscopy imaging

Time-lapse images were captured and analyzed by a confocal microscope LSM980 with ZEN software (ZEISS, Germany). Acquired data were annotated and analyzed using the ImageJ software. Images of *in situ* hybridization samples were captured using a DP70 light microscope (Olympus, Japan). Samples were positioned on 1% low-melting agarose mount in Petri dish covered with PBS.

### 4.12 Statistical Analysis and Graphs

Kruskal-Wallis tests were performed on each data set followed by one-way ANOVA analysis with Tukey’s post hoc analysis or student T-test. All statistical analyses were performed using GraphPad prism 10.3.1.

## ACKNOWLEDGEMENTS

We thank Maiko Yokouchi and Yuka Ueda for technical assistance with zebrafish breeding and experiments. Confocal images were acquired in the Biological Imaging Research at the Life Science Collaboration Center of NAIST. We are grateful to this center for being very helpful with confocal microscopy, image acquisition, and analysis.

---

**Supplementary Figure 1.**
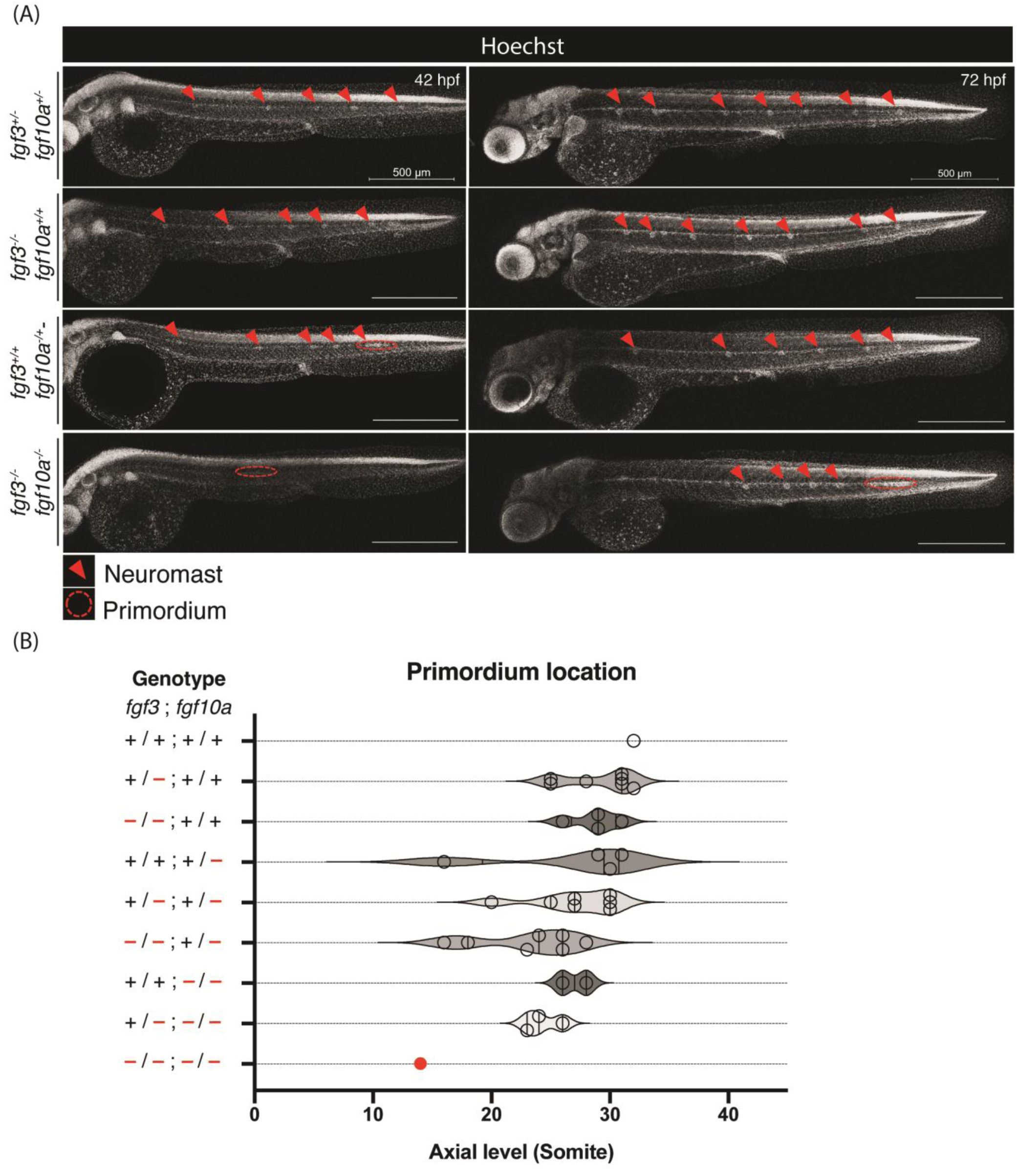
Posterior lateral line primordium and neuromast locations in early developing zebrafish embryos. (A) Confocal microscopic images of neuromast and primordium nuclei stained with Hoechst in *fgf3*,*fgf10a* heterozygous, single and double mutants at 42 and 72 dpf. (B) Schematic diagram showing axial somite-specific locations of posterior lateral line primordium nuclei in all genotypes at 42 hpf. Each dot represents one embryo.

**Supplementary Figure 2.**
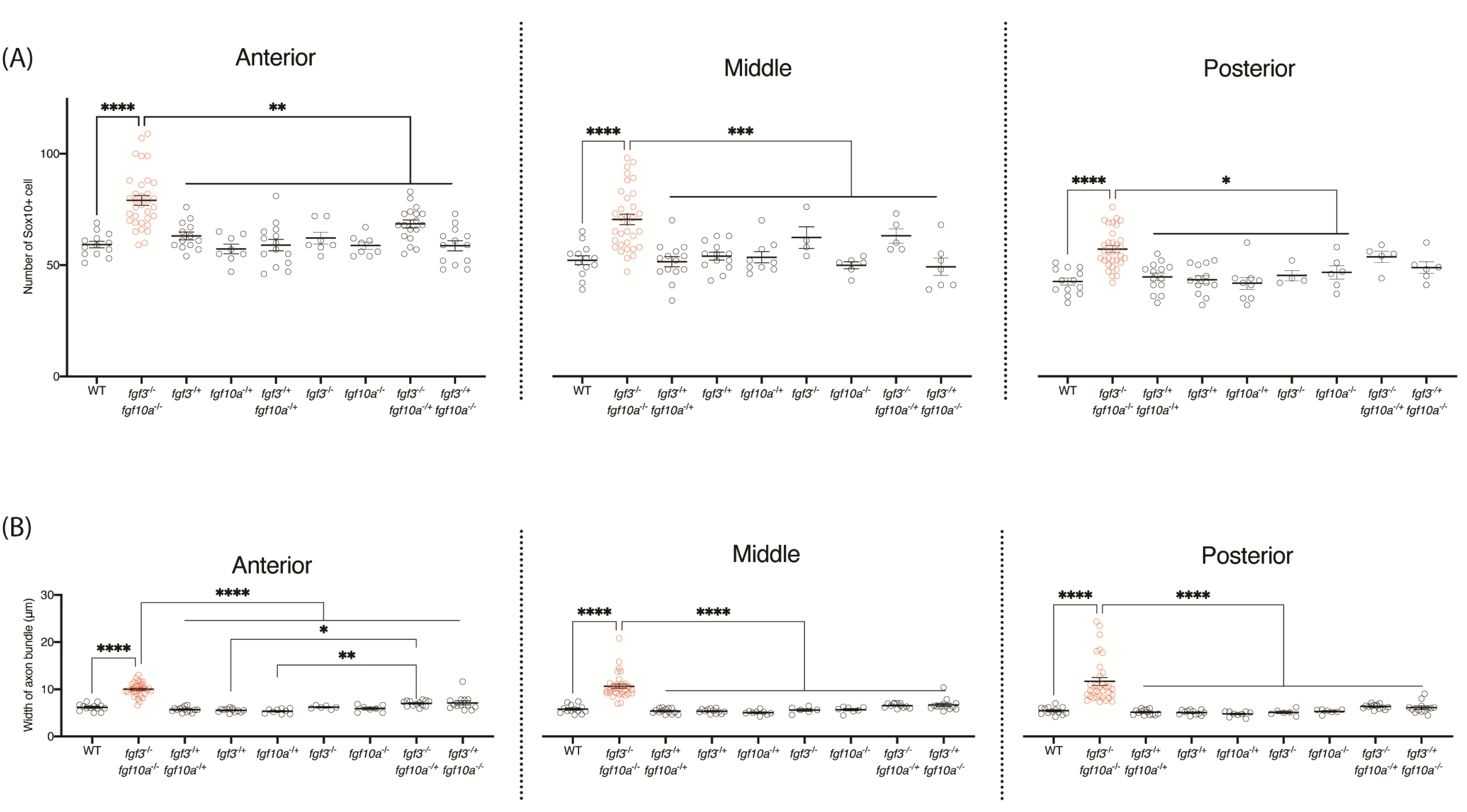
Quantification of mean number of So×10^+^ cell and the mean width of axon bundle. (A) the mean number of So×10^+^ cell and (B) the mean width of axon bundle in anterior, middle, and posterior regions in wildtype (WT) and *fgf3*,*fgf10a* double mutants and other genotype siblings at 5 dpf. Each dot represents a single larva. Data represent mean ± S.E.M and were analysed by one-way ANOVA with Tukey’s post-hoc multiple comparisons,* *p* < 0.05; ** *p* < 0.01; *** *p* < 0.005; **** *p* < 0.001.

**Supplementary Figure 3.**
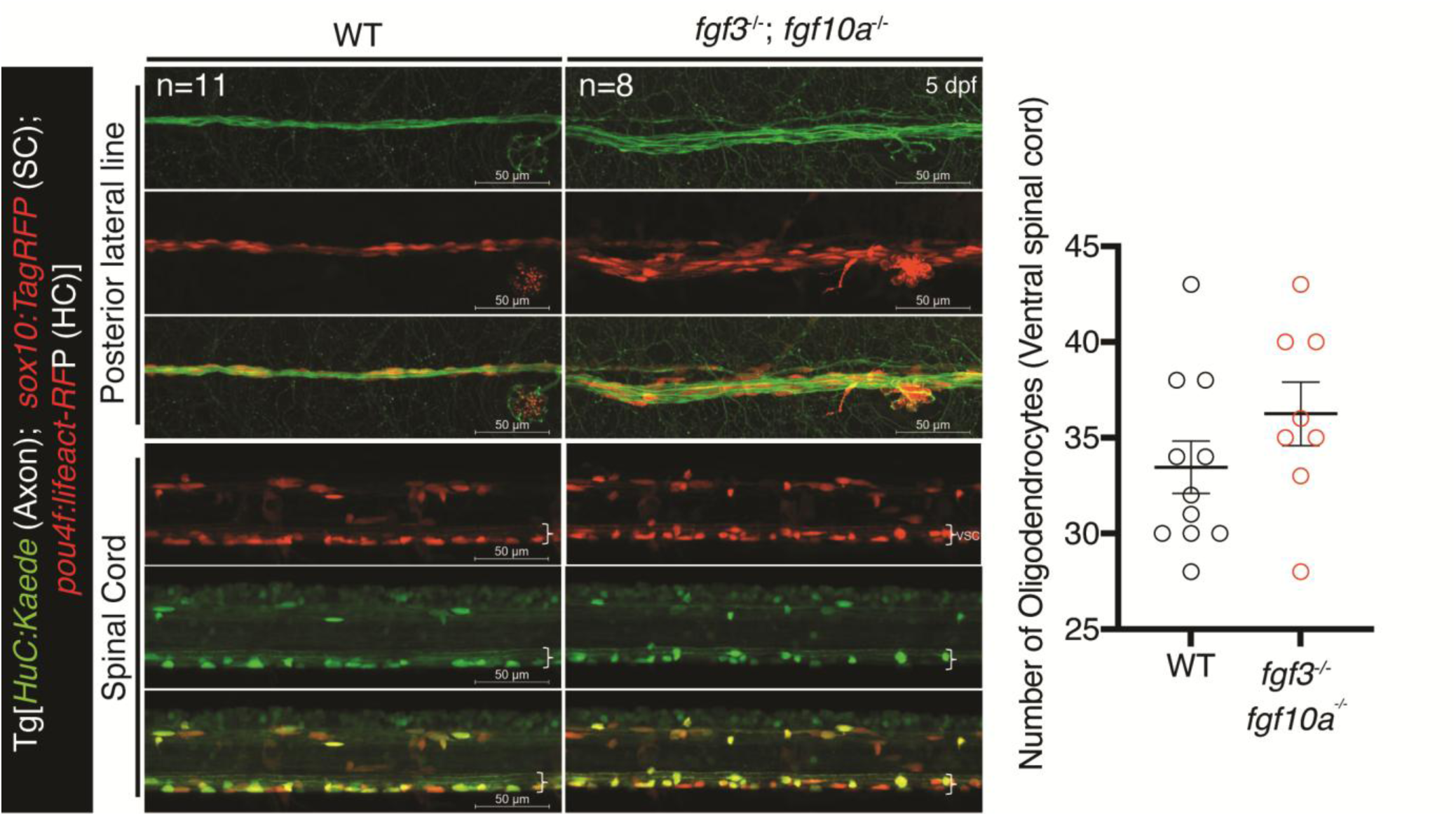
Schwann cell and axons in lateral line and spinal cord. Confocal live *in vivo* images of Schwann cell and axons, using Tg[*HuC:Kaede*;*zsox10:TagRFP*;*pou4f:lifeact-RFP*] in the middle region of posterior lateral line and spinal cord (15-17th somite) in wildtype (WT) and *fgf3*,*fgf10a* double mutants at 5 dpf. Scale bars: 50 μm. The quantification of oligodendrocytes in ventral spinal cord is also displayed in graph on the right. SC, Schwann cell, HC, hair cell; vsc, ventral spinal cord.

**Supplementary Figure 4.**
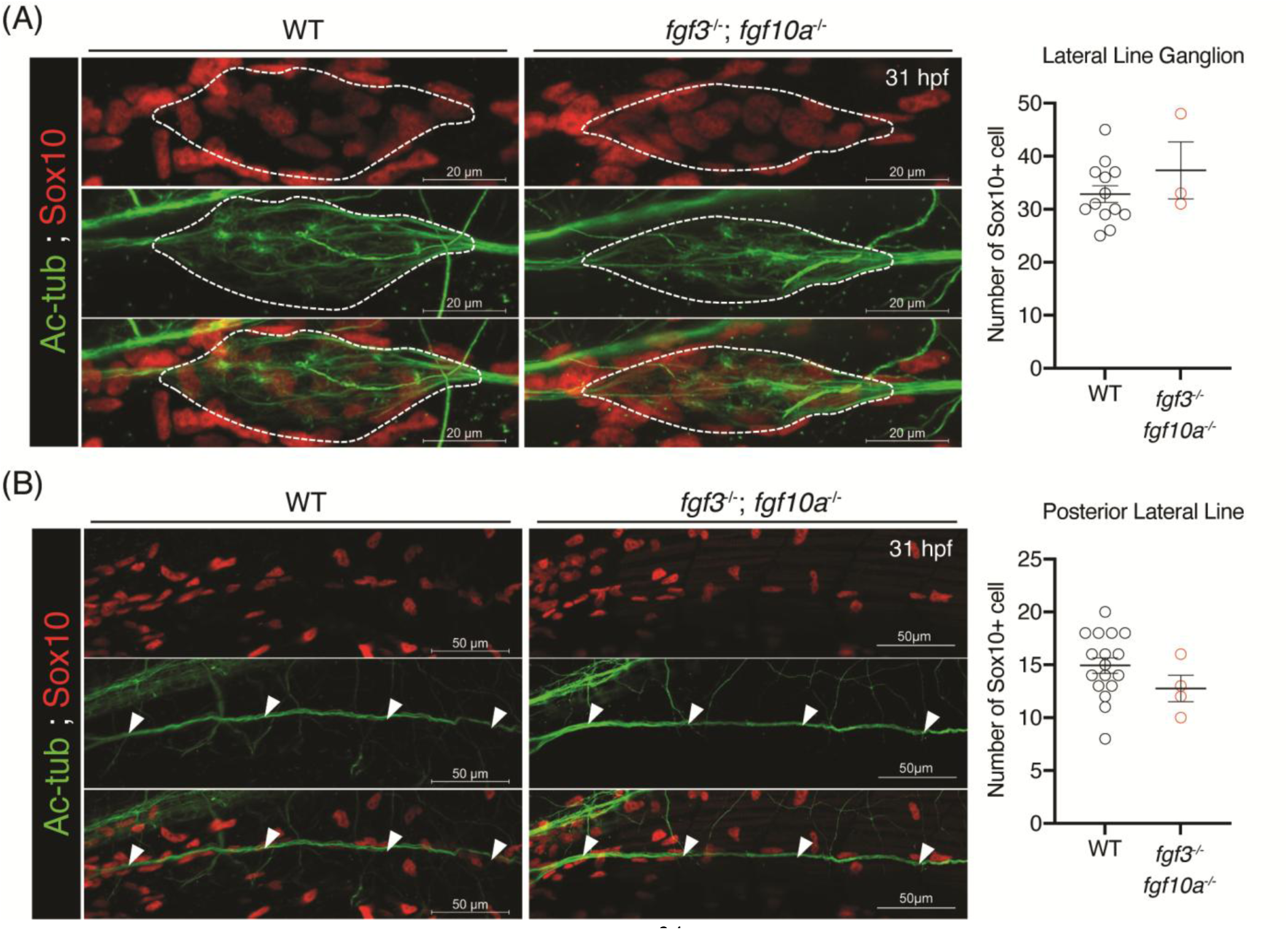
Confocal images of immunofluorescence staining visualizing the Schwann cells and axons in wildtype and *fgf3fgf10a* double mutants at 31 hpf. (A) Confocal images of So×10^+^ cells and axons labelled with anti-Sox10 and anti-Acetylated tubulin antibodies in pll ganglion and quantification of the mean So×10^+^ cell number in pll ganglion (B). Pll ganglion is indicated by the white dotted line. Anterior is to the left. Scale bars: 20 μm. (B) Confocal images of Sox10+ cells and axons labelled with anti-Sox10 and anti-Acetylated tubulin antibodies in pll axons and quantification of the mean Sox10+ cell number. Pll axons are indicated by white arrowheads. Anterior is to the left. Scale bars: 50 μm. Data represent mean ± S.E.M and were analysed by one-way ANOVA analysis with Tukey’s multiple comparisons (B) and student T-test (D).

**Supplementary Figure 5.**
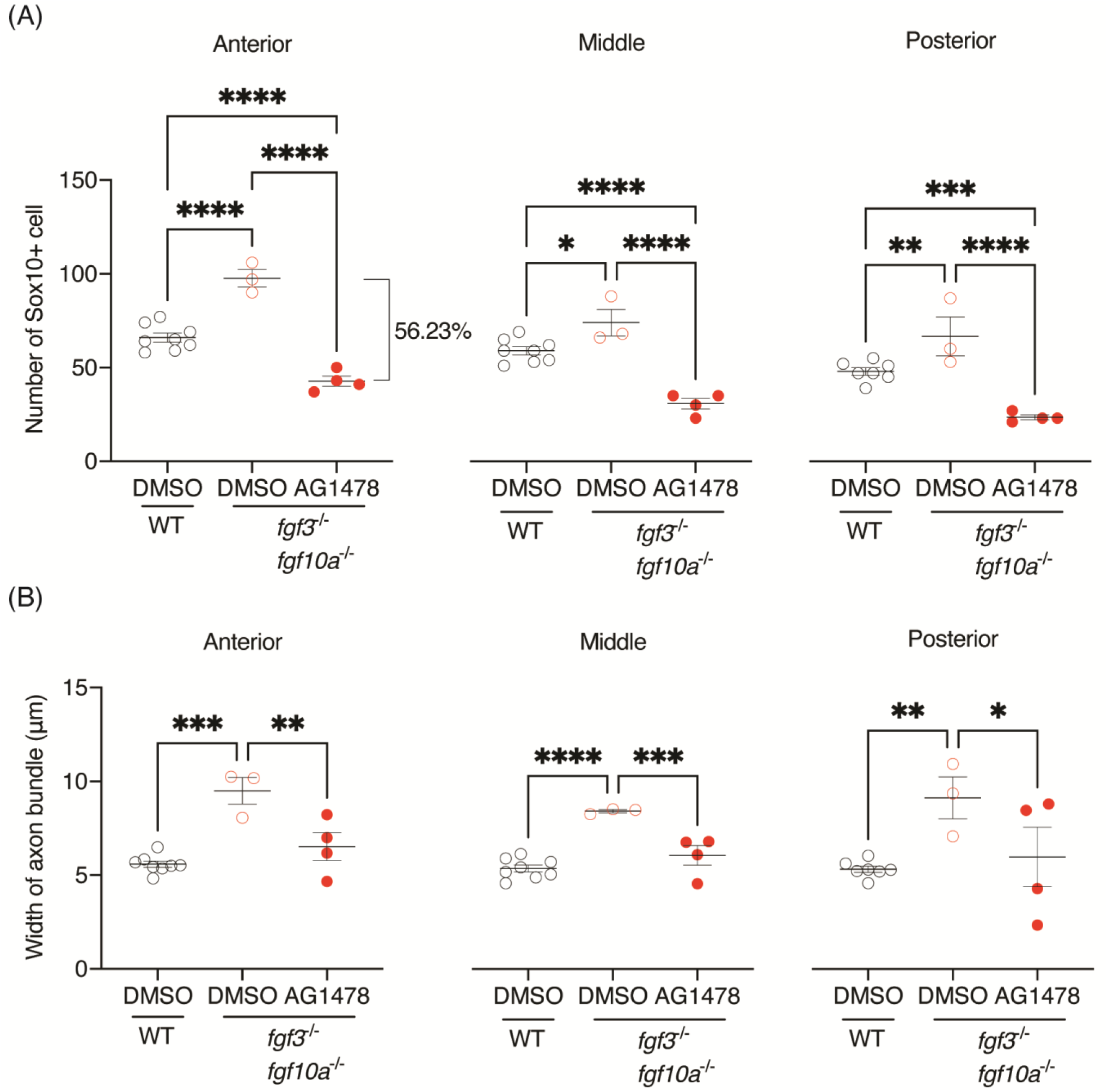
Quantification of So×10^+^ cell number and axon bundle width in AG1478 treated larva. Quantification of (A) the mean number of So×10^+^ cell and (B) the mean width of axon bundle in anterior, middle, and posterior regions in 2-5 dpf DMSO treated wildtype (WT), DMSO and AG1478 treated *fgf3*,*fgf10a* double mutants at 5 dpf. Each dot represents a single larva. Data represent mean ±S.E.M and were analysed by one-way ANOVA with Tukey’s post-hoc multiple comparisons,* *p* < 0.05; ** *p* < 0.01; *** *p* < 0.005; **** *p* < 0.001.

**Supplementary Movie 1.** Time-lapse imaging of a wild-type (WT) larva showing lateral line axons (Tg[HuC:Kaede]) and Schwann cells (Tg[sox10:TagRFP]) at 68–75 hpf.

Imaging was performed in Tg[*HuC:Kaede*; *sox10:TagRFP*; *pou4f:lifeact-RFP*]. Schwann cell proliferation and infiltration into interaxonal spaces are rarely observed, and axonal bundles remain tightly fasciculated. Blue arrowheads indicate dividing Schwann cells. This movie is representative of three independent larvae (n = 3).

**Supplementary Movie 2**. Time-lapse imaging of an *fgf3,10a* double mutant larva showing lateral line axons (Tg[HuC:Kaede]) and Schwann cells (Tg[sox10:TagRFP]) at 68–75 hpf.

Imaging was performed in Tg[*HuC:Kaede*; *sox10:TagRFP*; *pou4f:lifeact-RFP*]. Increased Schwann cell proliferation and frequent migration of daughter cells into interaxonal spaces are observed, accompanied by expansion of interaxonal space and axonal defasciculation. Blue arrowheads indicate dividing Schwann cells. Yellow arrowhead indicates daughter Schwann cells infiltrating into interaxonal spaces following cell division. This movie is representative of three independent larvae (n = 3).

**Supplementary Table 1.**
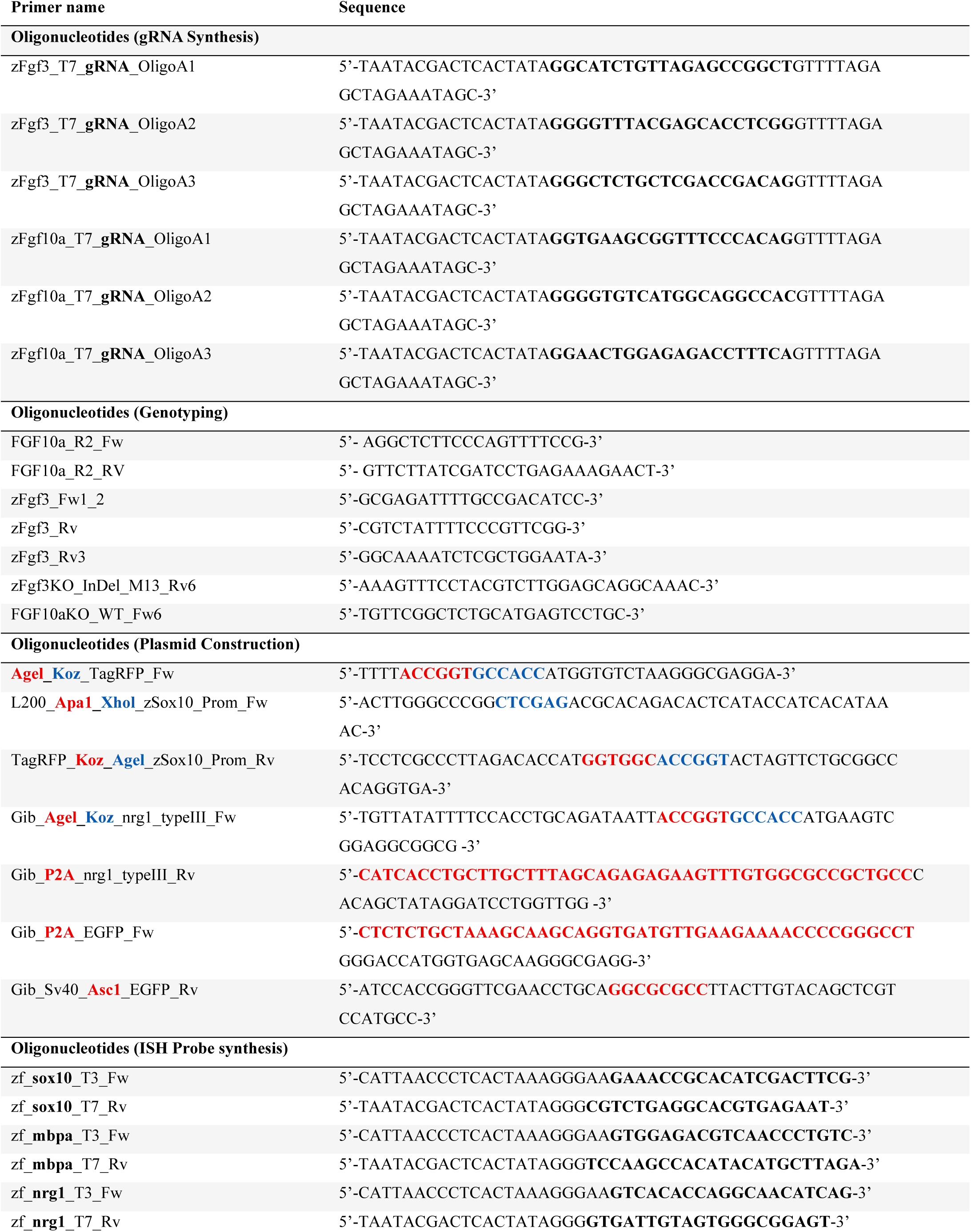
Primers and their sequences used in current study.

**Supplementary Table 2.** Primer pairs for PCR genotyping and its annealing temperatures.

| Genotype | Primer | Annealing temperature |
| --- | --- | --- |
| *fgf3<sup>WT</sup>* | zFgf3_Fw1_2 | 66.5°C |
|  | zFgf3_Rv3 |  |
| *fgf3<sup>13del</sup>* | zFgf3_Fw1_2 | 65°C |
|  | zFgf3KO_InDel_M13_Rv6 |  |
| *fgf10a<sup>WT</sup>* | FGF10aKO_WT_Fw6 | 65°C |
|  | FGF10a_R2_RV |  |
| *fgf10a<sup>7del</sup>* | FGF10aKO_InDel_M11_Fw3 | 65°C |
|  | FGF10a_R2_RV |  |

